# *Genome sequencing, assembly and annotation of the carob tree, Ceratonia siliqua* (Eudicots:Fabaceae)

**DOI:** 10.1101/2023.02.03.526947

**Authors:** Androniki C. Bibi, Panagiotis Ioannidis, Christos Bazakos, Kriton Kalantidis

## Abstract

The carob tree, *Ceratonia siliqua*, is an evergreen tree that belongs to the family of Fabaceae. It has been cultivated for thousands of years in Mediterranean countries and thus, has a considerable impact on the nutrition of humans as well as other animals of this region. Despite its importance, genomic resources are still scarce for this plant species. To fill this gap, we undertook the sequencing, assembly and annotation of the carob tree genome, which resulted in the first, nearly chromosome-level assembly for this plant species. The total assembly size is 492 Mbp which is close to the previously estimated genome size. The assembly N50 is 34.99 Mbp, thus showing a high contiguity, which is also evident from the fact that >98% of the assembly is contained in as few as 17 very large contigs. Moreover, both the genome sequence as well as the predicted gene set contain more than 96% of conserved, full-length BUSCOs. Finally, a comparative orthology analysis with 10 other plant species showed that the vast majority of the predicted carob tree genes have orthologs in the other plants and only a small fraction of them appears to be species-specific. Additional analyses are in progress for a more detailed study of the agronomic traits of this important legume.

## Introduction

The carob tree (*Ceratonia siliqua* L.) is an evergreen tree that belongs to the legume (Fabaceae) family. It has slow growth, great longevity and an extended flowering period (Diamantoglou and Mitrakos 1981). It originates from West Asia and is cultivated for years in Mediterranean countries (Santonocito 2020). Ancient Greeks brought it to Greece and Italy, while Arabs disseminated it along the North African coast and north into Spain and Portugal (Batlle and Tous 1997; Azab 2017). The carob tree is a very important part of the Mediterranean vegetation. It has been grown in mild and dry places with poor soils since antiquity, in most countries of the Mediterranean basin. It is also resistant to drought and salinity (Lo Gullo and Rosso 1986).

Carob pod production in the world is estimated to be about 310,000 tons per year and the principal producing countries are Spain, Italy, Portugal, Morocco, Greece, Cyprus, Algeria and Tunisia (Battle and Tous 1997; FAO 2022; Rtibia et al. 2017). Carob fruits consist of the pods as well as the seeds found inside the pods. They are considered powerful antioxidants due to the high content of bioactive phytochemicals. Carob fruit is enriched with several primary metabolite classes, such as sugars viz., D-pinitol “sugar alcohol” and a galactomannan gum, aside from proteins, fatty acids, minerals and dietary fibers. The carob secondary metabolites have attracted attention for their use as a functional food, mostly for their abundance in phenolic acids, tannins and flavonoids (Rasheed et al. 2017).

Carob trees are mostly dioecious but they can also be monoecious (Linskens and Scholten 1980; Batlle and Tous 1988). The genomic resources for the carob tree is scarce, thus limiting the understanding of the molecular basis of such basic biological features. Moreover, due to this paucity of genomic information, there are only speculations regarding the genome size of *C. siliqua* (Bureš et al. 2004). It is reported that the number of chromosomes of carob are 2n=24 (de Almeida, 1948; Frahm-Leliveld, 1957; Berger et al., 1958; Goldblatt, 1981; Yeh et al., 1986; Arista and Talavera, 1990; Rice et al. 2015; Peruzzi et al. 2016). It is also known that carob tree is diploid, even though some cases of triploidy and tetraploidy have been found (Bureš et al. 2004). Lately, based on genotypes from 17 microsatellite loci (including SSRs and SNPs) and geographical coordinates, 1,019 carob trees were grouped into seven carob evolutionary units (CEUs), with an interesting group being the E2, which involves the carob trees from Lebanon and Crete (Greece) (Figure 1) (Baumel et al. 2022). It appears that there is a high frequency of carob trees in the landscape of Lebanon and Crete. These are mostly found in abandoned fields or orchards in Lebanon, but cultivated and exploited in Crete.

**Figure 1.**
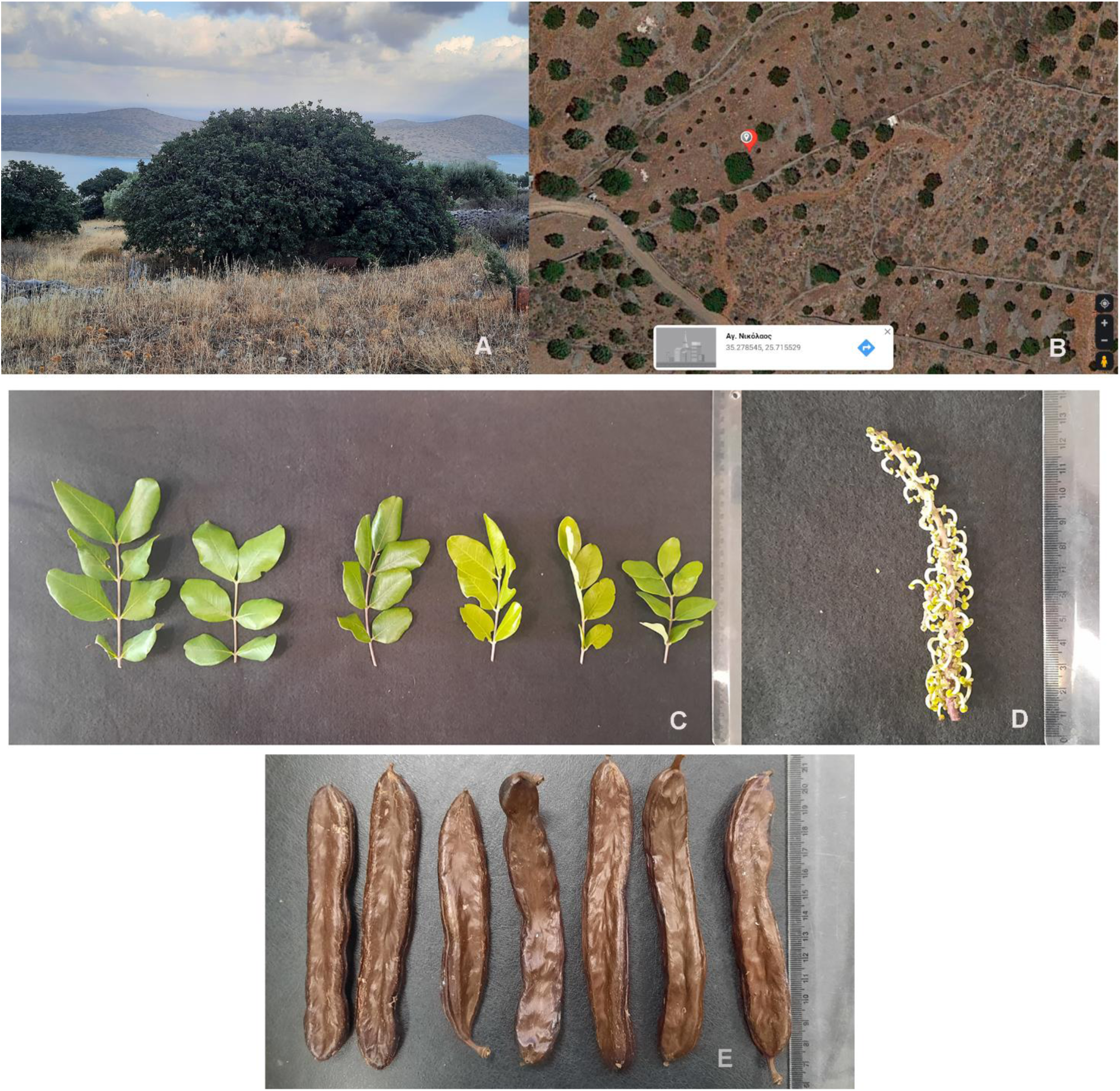
The Carob tree (A), the coordinates (B), various sizes of leaves (C), the female inflorescence (D), and the carob pods produced from this Carob tree (E).

Exploring worldwide biodiversity at the molecular level is of increasing interest during the last few years. This trend has resulted in the formation of national (e.g. the Darwin Tree of Life in Britain) and international (e.g. the European Reference Genome Atlas, or the Earth BioGenome Project) consortia that aim at sequencing all plants and animals on the planet. Therefore, species of regional importance, must be prioritized in genome sequencing efforts.

With these points in mind, we undertook the sequencing, assembly and annotation of the carob tree genome using a cultivar that is indigenous in Crete, Greece. The resulting, high quality genome is the first for this important plant species and was assembled using long read sequencing technology. Gene annotation was performed using standard Illumina RNAseq data as transcription evidence. The resulting gene set was used in a comparative analysis with other closely, as well as more distantly, related plant species and will shed light on key features of this interesting plant species. Among them, the most interesting ones are adaptation to dry climatic conditions, metabolic pathways related to the synthesis of valuable nutrients, and establishment of symbiotic relationships with nitrogen-fixing soil bacteria.

## Methods

### Plant material

A Cretan carob tree with specific qualitative and quantitative characteristics was chosen to be sequenced, in collaboration with the EPIMENIDIS cultural club of Panormos. This tree is located in Pines, Lasithiou in Crete (coordinates: L:35,278545, A:25,715529). Leaf tissue of an annual vegetation was collected and stored in −80 ultrafreezer for further analysis.

### Morphology

The selected carob tree has a stature that spreads about 4-5 m high, with branches spreading out to the ground (Fig. 1A). It is an evergreen plant with compound leaves, with 6-8 opposite leaflets light to dark green. Also this carob tree is monoecious, with female inflorescence, only with pistils.

### DNA extraction

Leaves of annual vegetation were ground with mortar and pestle in liquid nitrogen. DNA was extracted with a CTAB-based protocol. A CTAB buffer was prepared with 20.5g NaCl, 5g CTAB in 215mL ddH20, 25 mL 1M TRIS-HCL pH:8, 10 mL 0,5M EDTA pH:8, 1μL b-mercaptoethanol and 1% (w/v) PVP 360. An amount of 0.5 ml of CTAB buffer was added to the 50mg (0,05g) of ground leaf tissue and then incubated at 65°C for 30 min with occasional vigorous shaking. 0.5mL of a 24:1 chloroform: Isoamyl alcohol solution were added to each sample and shaked very well by vortex. The samples were centrifuged at 13,000 rpm for 15 min to resolve phases. The aqueous phase was pipetted out carefully to a fresh tube (1.5mL), added 0.5 mL cold isopropanol, mixed and incubated at −20°C or at 4°C or on ice for 1 hour or overnight. Subsequently, the samples were centrifuged at 9,000 rpm for 5 min. The supernatant was discarded and the pellet was washed in 70% ethanol. Following that, 1,000μL 70% ethanol was added and the samples were centrifuged at 13,000 rpm for 5 min and then the ethanol was carefully removed and drained dry (for at least an hour). The precipitate was dissolved in 100 μL TE buffer by gentle inversion and let it stay for at least an hour. We added 1μL RNase A (10mg/mL), incubated at 37°C for 30 min, RT for 15 min and then added 1μL Proteinase K (1mg/mL) and incubated at 37°C for 15-30 min. We then added 10μL of 3M sodium acetate and 250 μL absolute ethanol. We incubated the mixture at −20°C or at 4°C or on ice for 1 hour or overnight. The samples were centrifuged at 14,000 rpm for 10 min and the supernatant was discarded. Following that, 1ml of 70% ethanol were added and then centrifuged at 13,000 rpm for 5 min, carefully removed the ethanol and the samples were dried (for at least an hour). In the final step, the DNA pellets were suspended in 100 μL 1X Tris-EDTA buffer and stored at −20°C.

### RNA extraction

Newly developed leaves and pistils were collected from the same carob tree (coordinates: L:35,278545, A:25,715529). The tissues were stored in −80 °C ultra-freezer. Liquid nitrogen in a mortar and pestle was used to thoroughly grind the plant tissues. The Sigma Spectrum plant total RNA kit (Merck KGaA, Darmstadt, Germany) was used for RNA extraction that followed the manufacturer procedure. The degradation of RNA was observed with a denaturing gel and was quantified by Nanodrop and Qubit. The RNA integrity number for the samples was evaluated with Bioanalyzer RNA 6000 Nano / Pico Chip (Agilent). Two libraries were sent to Azenta for sequencing with the Illumina Novaseq 2×150 platform. A total of 32.2 and 35.2 million reads were generated from each library. These RNAseq reads are available from the Sequence Read Archive (SRA) under the BioProject accession PRJNA927143.

### Genome sequencing

Four PacBio libraries were sequenced, yielding a total of >10 million HiFi long reads, with a total length of >52 Gbp (Table 1). Since the estimated genome size for the carob tree is approximately 570 Mbp (Bures et al. 2004), our sequencing resulted in a sequencing depth of >90x, which is more than what is typically required for a decent assembly, based on PacBio reads. The genome sequencing reads are available from the Sequence Read Archive (SRA) under the BioProject accession PRJNA927143.

**Table 1.** Sequenced PacBio libraries.

| Library name | Number of reads | Total size (bp) | Average read length (bp) |
| --- | --- | --- | --- |
| Carob_1 | 2,575,892 | 13,207,640,173 | 5,127 |
| Carob_2 | 2,588,650 | 12,315,261,932 | 4,757 |
| Carob_3 | 2,572,906 | 13,847,707,024 | 5,382 |
| Carob_4 | 2,487,421 | 13,297,119,209 | 5,345 |
| **Total** | **10,224,869** | **52,667,728,338** |  |

### Genome assembly

The Hifiasm 0.16.1-r375 (Cheng et al 2021) assembler was used with only the “--primary” parameter and everything else set to the default values. Finally, duplicate contigs were removed using Purge_Dups v1.2.5 (Guan et al 2020), thus generating the final genome assembly (Table 2). *De novo* identification of species-specific repeat elements was carried out with RepeatModeler v1.0.11 (Smit and Hubley, repeatmasker.org/RepeatModeler), with parameters “-engine ncbi”. Subsequently, conserved as well as species-specific repeats were masked in the genome assembly using RepeatMasker v4.0.9 (Smit and Hubley, repeatmasker.org/RepeatMasker) with parameters “-xsmall -lib species-specific_repeats.fasta” (Table 3). The quality of the resulting assembly was assessed at the different points along the genome assembly process using BUSCO v3.0.2 (Waterhouse et al. 2018).

**Table 2.** Genome assembly statistics.

|  |  |
| --- | --- |
| **Size of PacBio reads** | 52.67 Gbp |
| **Assembly size** | 493 Mbp |
| **Sequencing coverage** | 107x |
| **Total number of contigs** | 118 |
| **Contig N50** | 34.99 Mbp |
| **GC content** | 33.66% |
| **BUSCO score (original assembly)**<br>(Embryophyta, 1,375 BUSCOs) | Complete: 96.2% |
|  | of which duplicated: 8.1% |
|  | Fragmented: 1.2% |
|  | Missing: 2.6% |
| **BUSCO score (purged assembly)**<br>(Embryophyta, 1,375 BUSCOs) | Complete: 96.1% |
|  | of which duplicated: 8.1% |
|  | Fragmented: 1.2% |
|  | Missing: 2.7% |
| **Number of predicted genes** | 43,965 |

**Table 3.** Classes of identified repeat elements.

| Repeat class | Length occupied | Percentage of total genome sequence |
| --- | --- | --- |
| SINEs | 75,890 | 0.02 |
| LINEs | 2,002,509 | 0.41 |
| LTRs | 83,037,038 | 16.84 |
| DNA elements | 12,770,676 | 2.59 |
| Small RNA | 1,604,765 | 0.33 |
| Satellites | 39,103 | 0.01 |
| Simple repeats | 5,910,529 | 1.2 |
| Low complexity | 1,540,842 | 0.31 |
| Unclassified, interspersed | 112,804,239 | 22.87 |
| Unclassified, others | 939,436 | 0.19 |
| **Total** | **220,725,027** | **44.76** |

### Gene prediction

The BRAKER pipeline v2.1.3 (Bruna et al 2021) was used in ETP mode with parameters “--gff3 –prg=gth –cores=16”. Two RNAseq Illumina libraries from *C. siliqua* were used as transcription evidence with the “--bam=” parameter. These libraries contained pooled RNA from leaves, stamens and pistils. Additionally, we used RNAseq data from another three related Fabaceae species; *Senna tora, Acrocarpus fraxinifolius* and *Acacia pycnantha*. Finally, the complete gene sets of closely related Fabaceae species were passed to BRAKER as protein evidence with the “--prot_seq” parameter. In order to reduce redundancy, we removed 100% identical proteins in the above set of proteins, using CD-HIT v4.7 (Fu et al 2012) with parameters “-n 5 -c 1.00 -M 64000”. The final gene set contained 43,965 genes (Table 4) and was used in a comparative genomics analysis based on Orthofinder v2.5.4 (Emms and Kelly 2019), with another ten plant species (Table 5).

**Table 4.** Features of the predicted gene set.

|  |  |
| --- | --- |
| Number of proteins (including isoforms) | 47,133 |
| Number of genes (without isoforms) | 43,965 |
| ...with a significant hit in SwissProt | 25,651 |
| ...with a significant hit in Uniref50 | 34,470 |
| ...with a Pfam domain | 26,370 |
| ...with an assigned GO term | 17,240 |
| ...assigned to a PANTHER pathway | 27,685 |
| **BUSCO score (gene set w/o isoforms)**<br>(Embryophyta, 1,375 BUSCOs) | Complete: 96.3% |
|  | of which duplicated: 8.8% |
|  | Fragmented: 2.2% |
|  | Missing: 1.5% |

**Table 5.** Plant species used for the comparative genomics analysis.

| **Species** | **Common name** | **Family** | **Number of genes** | **Accession** |
| --- | --- | --- | --- | --- |
| *Arabidopsis thaliana* | Thale cress | Brassicaceae | 27,529 | GCF_000001735.4 |
| *Glycine max* | Soybean | Fabaceae | 46,894 | GCF_000004515.6 |
| *Malus domestica* | Apple tree | Rosaceae | 35,893 | GCF_002114115.1 |
| *Medicago truncatula* | Barrelclover | Fabaceae | 31,820 | GCF_003473485.1 |
| *Oryza sativa* | Rice | Poaceae | 28,685 | GCF_001433935.1 |
| *Phaseolus vulgaris* | Bean | Fabaceae | 28,134 | GCF_000499845.1 |
| *Pisum sativum* | Pea | Fabaceae | 40,025 | GCF_024323335.1 |
| *Solanum lycopersicum* | Tomato | Solanaceae | 25,431 | GCF_000188115.5 |
| *Triticum aestivum* | Wheat | Poaceae | 103,751 | GCF_018294505.1 |
| *Zea mays* | Maize | Poaceae | 34,174 | GCF_902167145.1 |
| *Ceratonia siliqua* | Carob | Fabaceae | 43,965 | This study |

## Results and discussion

### Genome assembly overview

The genome of the carob tree, *C. siliqua*, was assembled using >10 million long, high quality (HiFi) PacBio reads. We used the Hifiasm assembler to obtain a highly contiguous assembly comprising 1,402 contigs with a total size of 518 Mbp and contig N50 of 34.99 Mbp. Purging duplicate contigs with the Purge_Dups program resulted in an assembly containing 118 contigs and a total size of 492 Mbp, while the contig N50 remained the same. The gene content of the assembly was assessed with BUSCO (Waterhouse et al. 2018) and found to be excellent, since it contains the complete sequence of 96.1% of the 1,375 Embryophyta BUSCOs (Figure 2). Purging did not affect the BUSCO scores, which suggests that the majority of the purged contigs were non-coding. The high quality of the genome assembly is indicated by the fact that the vast majority of the sequence (485 Mbp of a total 492 Mbp) in the purged assembly is found in only 17 contigs with a length >10 Mbp.

**Figure 2.**
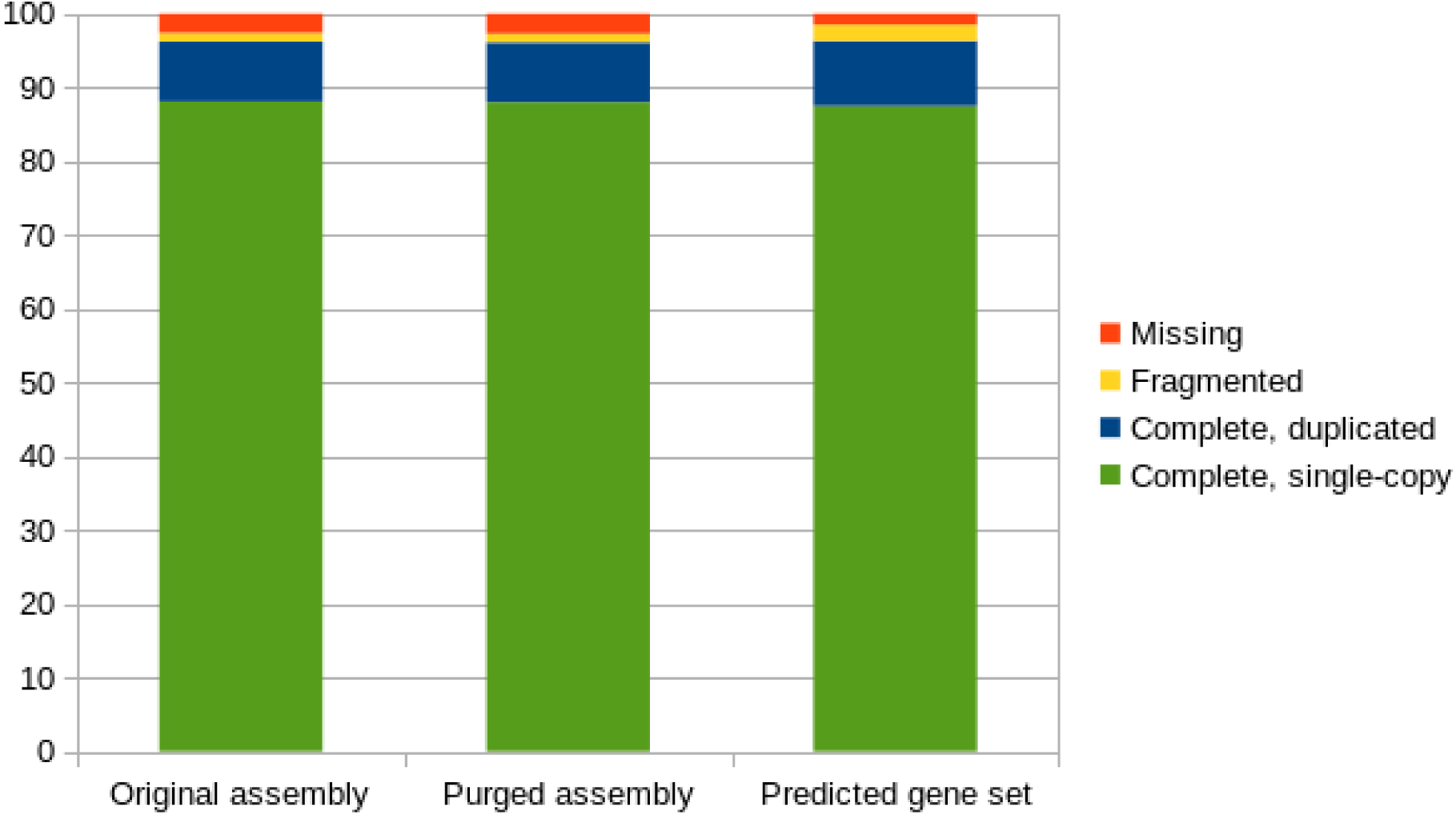
Comparative BUSCO scores (original, purged and gene set).

### Predicted gene set

Different classes of repeated DNA elements were identified, most notably LTRs (long terminal repeat elements) covering approximately 80 Mbp (15.5%) in the final genome assembly (Table 3). The BRAKER pipeline predicted 43,965 protein-coding genes, a number which is comparable to that of other plant species (Table 5). Most importantly, the BUSCO score of the predicted gene set was better (96.3% complete BUSCOs) than that of the genome assembly (96.1%). An additional 2.2% of BUSCOs were found fragmented in the gene set and only 1.5% of BUSCOs were missing. Overall, the BUSCO scores for the predicted gene set are better compared to those from the genome assembly, indicating that the gene prediction pipeline worked very efficiently and did not miss conserved genes that exist in the assembly (Figure 2). Consequently, the resulting gene set is as complete as possible and thus suitable for the downstream analyses. Of the 43,965 genes, 34,735 have at least one of the following: (a) a significant (e-value < 1e-05) hit against the SwissProt database, (b) a significant hit against Uniref50, (c) a Pfam domain, (d) have been assigned a Gene Ontology (GO) term, (e) have been assigned to a PANTHER pathway (Table 4).

### Orthology analysis

An orthology analysis including ten more plant species (Table 5) was undertaken in order to study gene family evolution in those species. A species tree was reconstructed in Orthofinder using information from the individual gene trees (Figure 3A). The topology of the species tree is the expected one, with all Fabaceae (including *C. siliqua*) forming a monophyletic clade and the representatives from Rosaceae (*M. domestica*), Solanaceae (*S. lycopersicum*), and Brassicaceae (*A. thaliana*) forming sister clades to Fabaceae. Lastly, all three monocot species form a monophyletic clade that is sister to all previously mentioned dicots. Clustering the entire gene sets of these 11 species showed that the majority of the genes were assigned to orthogroups (Figure 3B). Further analysis, showed that *C. siliqua, T. aestivum* and *P. sativum* contain a relatively large fraction of species-specific genes (Figure 3C), which warrants further investigation.

**Figure 3.**
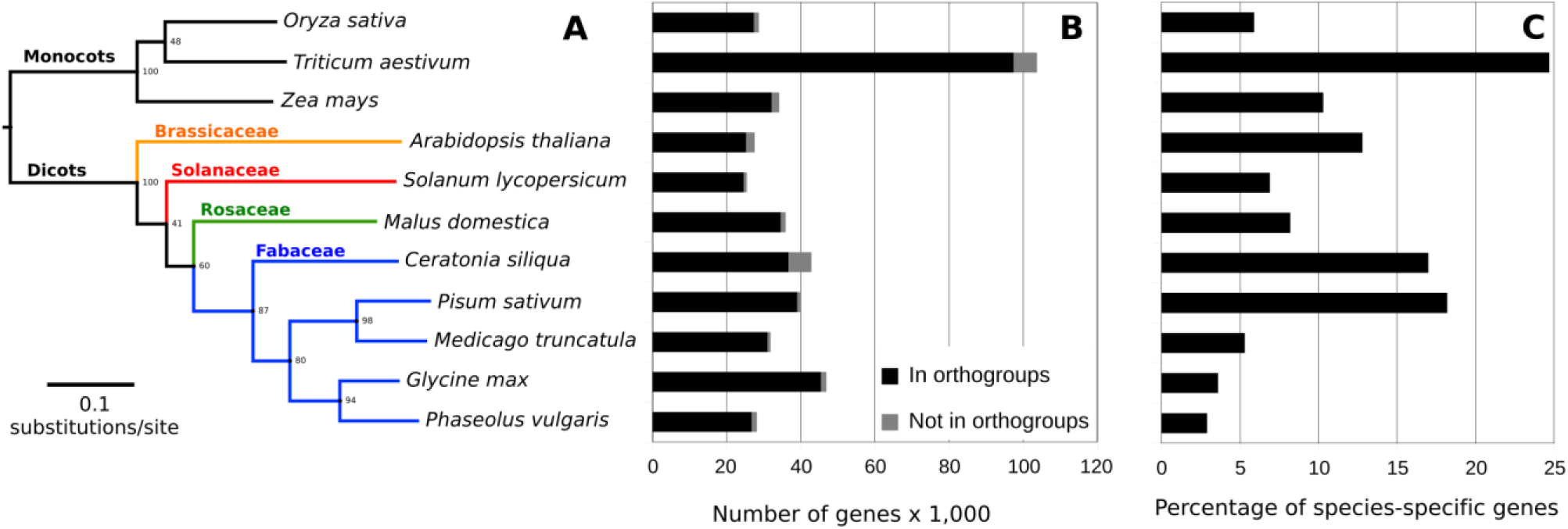
Species tree and orthology analysis. **(A)** Species tree including carob tree (*C. siliqua*) and ten more plant species. **(B)** Number of genes assigned to orthogroups (black), or not assigned in any orthogroup (grey). **(C)** Percentage of the total number of genes that is species-specific.

## Conclusions

The high quality genome assembly for the carob tree, a plant species of high regional importance for the Mediterranean countries, is reported in this study. The current genome assembly is nearly at chromosome-level, with virtually the entire assembly contained in only 17 very large (>10 Mbp) contigs. Moreover, the gene content is also very high as shown by the BUSCO metrics, whereby >96% of the Embryophyta set were found as complete, both in the genome assembly, as well as in the predicted gene set. This high quality assembly, paired with transcriptome data that were generated within the frame of this project, resulted in a very complete predicted gene set. The gene set was used in order to perform a brief comparative analysis in which carob was compared to ten other plant species. The resulting species tree is in agreement with the already established phylogeny, placing carob as basal to all other Fabaceae. Additional analyses are in progress in order to further improve the current genome assembly using Hi-C scaffolding. Subsequently, detailed comparative analyses will focus on key gene families implicated in biological processes that are characteristic for carob tree.

## Acknowledgements

This research has been financed by the Region of Crete through the “Actions for the utilization of the Carob tree potential (*Ceratonia siliqua*) in the Region of Crete” (project code: ADAM21SYMV008995374). Also, we would like to acknowledge Mrs Korina Miliaraki for her initiative in this project and continuous support and Mr. Drakonakis from Pines, Lasithiou for providing us the genetic material.

## Notes

Competing Interests: None

### Competing Interest Statement

The authors have declared no competing interest.

### Summary of Updates

We forgot to add the finance support in the acknowledgements

